# Interplay between positive and negative regulation by B3-type transcription factors is critical for the accurate expression of the *ABA INSENSITIVE 4* gene

**DOI:** 10.1101/2021.10.18.464865

**Authors:** Alma Fabiola Hernández-Bernal, Elizabeth Cordoba, Mónica Santos Mendoza, Kenny Alejandra Agreda-Laguna, Alejandra Dagmara Rivera, Maritere Uriostegui-Arcos, Mario Zurita, Patricia León

## Abstract

The ABA-INSENSITIVE 4 transcription factor is key for the regulation of diverse aspects of plant development and environmental responses, including proper perception of hormonal and nutritional signals. ABI4 activity is highly regulated at the transcriptional and post-transcriptional levels leading to precise expression mainly in the developing seed and early seedling development. Based on genetic and molecular approaches in the current study we provide new insights into the central mechanism underpinning the transcriptional regulation of *ABI4* during both seed and vegetative development. We identified a complex interplay between the LEC2 and ABI3 transcriptional activators and the HSI/VAL repressors that is critical for proper *ABI4* expression. Interestingly, the regulation by these proteins relies on the two RY *cis*-acting motifs present two kb upstream of the *ABI4* gene. Our analysis also shows that the chromatin landscape of the *ABI4* loci is highly dependent on the LEC2 and HSI2/VAL proteins. LEC2 regulation extends to the vegetative development and the absence of this factor results in ABA- and sugar-insensitive signaling in the developing plant. This regulatory circuit functions as a major control module for the correct spatial-temporal expression of *ABI4* and prevents its ectopic accumulation that is harmful to the plant.

## INTODUCTION

The transcriptional factor ABA-INSENSITIVE 4 (ABI4) is a member of the APETALA-2 (AP2)/ERF gene family, and is conserved in plants (Wind et al., 2013). ABI4 plays essential roles integrating nutritional, hormonal, abiotic, biotic and developmental signals (Chandrasekaran et al., 2020). However, ABI4 is a versatile regulator, required for abscisic acid (ABA) signaling and proper interaction with gibberellins (GA) and auxins during seed maturation, germination and post-germinative growth (Soderman et al., 2000; Shkolnik-Inbar and Bar-Zvi, 2010; Huang et al., 2017). ABI4 is also essential for sugar perception, nitrate sensitivity and ABA-dependent lipid mobilization in the embryo (Arenas-Huertero et al., 2000; Huijser et al., 2000; Rook et al., 2001; Signora et al., 2001; Penfield et al., 2006). Mutants of ABI4 are tolerant to salt (Quesada et al., 2000) and display defects in redox homeostasis (Kerchev et al., 2011). Finally, ABI4 regulate lateral root initiation (Shkolnik-Inbar and Bar-Zvi, 2010), male sterility and mitochondria- and chloroplast- retrograde communications (Koussevitzky et al., 2007; Giraud et al., 2009).

ABI4 affects the expression of diverse genes, acting as both a positive and a negative regulator by interacting with the CE1 (CACCG) sequence and related *cis-* acting elements (Niu et al., 2002; Acevedo-Hernandez et al., 2005; Koussevitzky et al., 2007; Wind et al., 2013). ABI4 induces the expression of genes such as the starch branching enzyme (*SBE2)* and the transcriptional factor *ABI5* in the presence of sugars (Bossi et al., 2009). In response to ABA, ABI4 also upregulates the expression of genes involved in ABA biosynthesis and GA catabolism, such as *NCED6* and *GA2ox7* (Shu et al., 2016b), in lipid catabolism, as oleosin and dehydrin (Penfield et al., 2006; Yang et al., 2011), and the flower transition gene *FLOWERING LOCUS C* (*FLC)* (Shu et al., 2016a) and *PHYTOCHROME A* (*PHYA)* (Barros-Galvao et al., 2020). In contrast to transcriptional activation, ABI4 represses the expression of diverse photosynthetic-related genes (PhANGS), the cytokinin response regulators (ARRs) and some genes involved in ethylene biosynthesis (Koussevitzky et al., 2007; Dong et al., 2016; Huang et al., 2016a). Recently, ABI4 has been shown to downregulate the *VTC2* gene that is required for plant defense responses (Yu et al., 2019).

The accumulation and activity of ABI4 is tightly controlled at the transcriptional and protein levels. At the protein level ABI4 is subjected to selective degradation via proteasome (Finkelstein et al., 2011; Gregorio et al., 2014) and its activity is modulated through by MAP kinase phosphorylation in response to sugars, ABA and salt stresses (Eisner et al., 2021). At the transcriptional level, *ABI4* is expressed predominately in the developing seed during germination and in the first days of the seedling development (Soderman et al., 2000; Penfield et al., 2006; Bossi et al., 2009). Later in development, the expression of *ABI4* is restricted to specific regions such as the vascular system of the petiole, pollen and the mature zone of the root (Shkolnik-Inbar and Bar-Zvi, 2010). Finally, the expression of *ABI4* is activated by environmental signals such as ABA and high sugar levels (Arroyo et al., 2003).

A central regulator of *ABI4* is the ABI4 protein itself, functioning as an activator to maintain its correct temporal-spatial transcription during early seedling development (Bossi et al., 2009). Other positive regulators that dictate the correct expression of *ABI4* in the germinating seed include the transcription factors MYB96, WRKY6 and the chloroplast envelope bound PTM (Sun et al., 2011; Lee et al., 2015; Huang et al., 2016b). The expression of *ABI4* is also downregulated by several factors including SCARECROW, WRKY18/40/60, RAV1 in the root apical meristem and BASS2 in the germinating seedlings (Shang et al., 2010; Cui et al., 2012; Feng et al., 2014; Zhao et al., 2016).

In spite that a major location of *ABI4* expression is in the developing embryo, the regulators responsible for its spatial and temporal expression remain elusive. The accumulation of ABI4 overlaps with that of the LAFL regulators of seed development, which include LEAFY COTYLEDON 1 (LEC1), LEC2, FUSCA3 (FUS3) and the ABA-INSENSITIVE 3 (ABI3) transcription factors (Le et al., 2010; Boulard et al., 2017; Lepiniec et al., 2018). LEC1 shares sequence similarity with the HAP3 subunit of the CCAAT-binding transcription factor and is a member of the NF-YB family (Lotan et al., 1998). In contrast, LEC2, ABI3 and FUS3 belong to the plant-specific B3-domain family (ALF), related to the maize VP1 protein (Stone et al., 2001). The LAFL regulators work in an intricate network and are essential for the regulation of key proteins required for the correct seed maturation and germination, embryonic identity, somatic embryogenesis, the acquisition of desiccation tolerance and dormancy, specification of the cotyledon identity and hormone signaling (Meinke et al., 1994; Santos-Mendoza et al., 2008; Tao et al., 2017; Lepiniec et al., 2018; Wang et al., 2020). Accordingly, mutants of the LAFL genes display diverse homeotic alterations, such as the acquisition of vegetative characters in the embryonic tissues (presence of trichomes in cotyledons) and precocious germination (Meinke et al., 1994; Lotan et al., 1998).

The mechanism of action of these regulators is diverse. For example, LEC1 acts as a pioneer transcriptional regulator promoting an active chromatin state activating transcription of the *FLC* gene (Tao et al., 2017). In contrast, LEC2 and FUS3 have been shown to activate the expression of various genes by displacing negative regulators, such as the *HIGH LEVEL EXPRESSION OF SUGAR INDUCIBLE GENE 2/VIVIPAROUS1/ABI3-LIKE 1 (HSI2/VAL1)* or *HSI2-LIKE1/VAL2* (*HSL1/VAL2*) proteins, two members of the B3-type family that act as central repressors of diverse seed developmental genes. These proteins interact with the same *cis*-acting sequences as the ALF (Tsukagoshi et al., 2007; Tao et al., 2019).

Previous studies showed that genetic interactions between ABI3, LEC1 and FUS3 with ABI4 in responses to ABA, sugar perception and development (Soderman et al., 2000; Brocard-Gifford et al., 2003). However, the molecular nature underlying these interactions remains unclear, since yeast two-hybrid analysis did not show direct interaction, nor that the level of the *ABI4* transcript was significantly altered in the *lec1* or *fus3* mutant backgrounds (Soderman et al., 2000; Brocard-Gifford et al., 2003).

Due to the role of ABI4 as an integrator of diverse signals, understanding the mechanisms that regulate its accumulation under diverse developmental and environmental conditions is important. In the present study using genetic and molecular analyses we show that several of the LAFL transcription factors are required to maintain the level and the correct expression pattern of *ABI4*. We identify LEC2 and ABI3 as critical direct activators of *ABI4* expression during seed development and early vegetative growth. Furthermore, our analysis uncovered an unexpected function of the HSI2/VAL1 as a major repressor of *ABI4* expression in vegetative tissues. The interplay between activation and repression exerted by these regulators occurs through the same *cis*-acting sequences and is essential for the correct expression of *ABI4* and its response to environmental signals such as sugar and ABA levels.

## RESULTS

### ABI4 expresses during all stages of the developing seed

During seed development, the expression of *ABI4* is restricted to the embryo (Soderman et al., 2000; Bossi et al., 2009). To obtain a detailed picture of the *ABI4* expression profile at different stages of the developing seed, we analyzed the b-glucuronidase (GUS) activity of the p*ABI4*:GUS transgenic line containing 3Kb of the *ABI4* regulatory region, which was previously shown to accurate reflect the expression of the endogenous transcript (Bossi et al., 2009). As shown in Figure 1A, we confirmed that *ABI4* expression restricts to the embryo proper and is detected at the pre-globular stage and in all the following developmental stages, except for the dry seeds as previously reported (Bossi et al., 2009).

**Figure 1.**
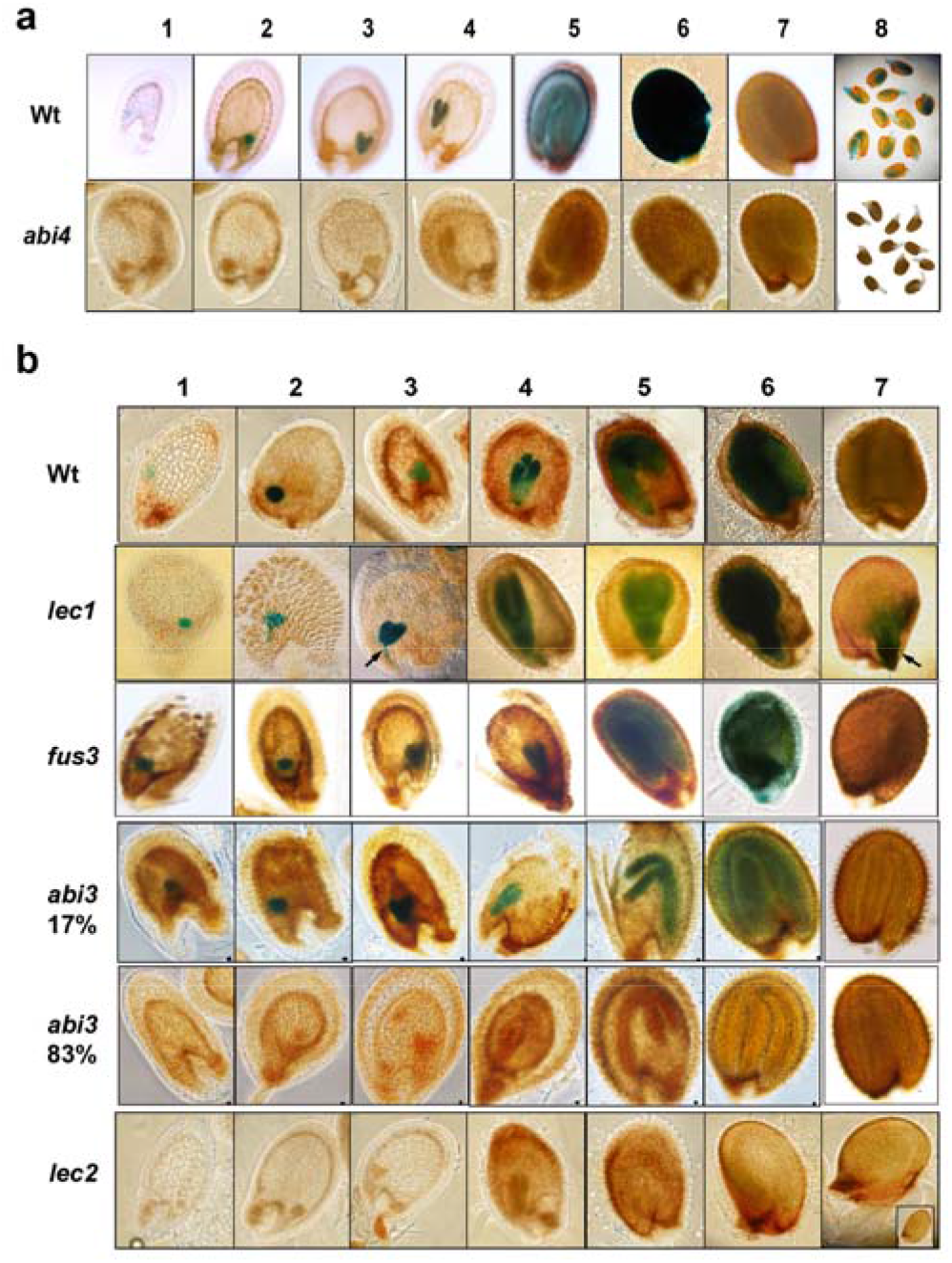
ABI4 and the LAFL transcription factors regulate the expression of *ABI*4 during seed development. A) Expression pattern of the 3kb *ABI4*:GUS transgene monitored in wild-type (Wt) and *abi4* mutant embryos at preglobular (1), globular (2), heart (3), torpedo (4), bent cotyledon (5), mature (6) dry seeds (7) and 24h germinating seedlings (8). A representative pattern for each line is shown. B) Pattern of GUS expression in Wt, *lec1, fus3, abi3* and *lec2* mutant embryos at preglobular (1), globular (2), heart (3), torpedo (4), bent cotyledon (5), mature (6) developing seeds and dry seeds (7). The different expression patterns observed in the *abi3* mutant are shown and the percentage (%) of each is included. Arrow points to the suspensor tissue in the *lec1* mutant

Previous research demonstrated that ABI4 is an essential activator of its own expression during germination and early seedling development (Bossi et al., 2009). In this study we confirmed that ABI4 is also required for its expression during seed development, as no GUS activity of the *pABI4*:GUS transgene was detected in the *abi4* mutant background (Figure 1A). This data further confirms the critical auto-activation function of ABI4.

### The LAFL transcription factors regulate *ABI4* expression

Given the similarities in the temporal expression between ABI4 and the LAFL regulators during seed development, we evaluated their impact on the expression of *ABI4*. Therefore, we introduced the *3Kb pABI4:GUS* transgene (*pABI4:*GUS) into the *lec1*, *lec2, fus3* and *abi3* mutant backgrounds and analyzed the GUS temporal and spatial expression throughout seed development. As shown in Figure 1B we did not detect any major differences of *ABI4* expression in the *lec1* or *fus3* mutants compared to wild-type seeds, except for an ectopic expression of *ABI4* in the suspensor tissue in the *lec1* mutant that is maintained even in the dry seeds (Figure 1B *lec1* panels 2-7). In the homozygous *abi3* mutant seeds we observed two contrasting expression patterns where 17% of the seeds display a pattern similar to wild-type, but in 83% of the seeds no GUS activity was detected in any stage of the developing seed (Figure 1B *abi3* panels). Remarkably, in the case of the *lec2* mutant, GUS activity was undetectable in all stages of the developing seeds compared to wild-type (Figure 1B *lec2* panels). We corroborated that the absence of GUS activity in the *abi3* and *lec2* mutant backgrounds was not caused by mutations or silencing of the *pABI4*:GUS transgene as this reporter accumulates at normal levels in the corresponding heterozygous mutant seeds (Figure S1). Altogether these results provide novel insights into the regulation of the *ABI4* gene during seed development, where the transcription factors ABI3 and, in particularly LEC2, play central roles as positive regulators, while LEC1 has a negative role restricted to the suspensor tissue.

### LEC2 but not ABI3 is essential for *ABI4* expression during early seedling development

Previous studies showed that the *ABI4* transcript accumulates during germination and early seedling development (Soderman et al., 2000; Arroyo et al., 2003; Bossi et al., 2009). Given that our previous analyses showed important alterations in the expression of *ABI4* in the *lec2* and *abi3* mutants during seed development, we were interested to determine whether these transcription factors affect the temporal and/or spatial expression of *ABI4* in germinating seedlings. Therefore, we analyzed the GUS activity of the *pABI4*:GUS transgene in germinating seedlings of *lec1, lec2* and *abi3*. Since mutants of the LAFL transcription factors are desiccation intolerant, we collected the homozygous *lec1*, *lec2* and *abi3* and wild-type seeds prior to desiccation and transferred them to media for germination. Expression of the GUS reporter was detected in the *lec1* mutant in more than 98% of the germinating seeds (Figure 2E) as in the 3 day-old seedlings (Figure 2F), displaying a similar pattern to the wild-type (Figure 2A and 2B). In the case of the *abi3* mutant we also detected GUS expression in 100% of the germinating seedlings (Figure 2C) that is maintained in the 3 day-old plants (Figure 2D), albeit at a lower level than in the wild-type seedlings (Figures 2A and 2B). These results support a role for ABI3 in the maintenance of the expression level of *ABI4* during early seedling development. Interestingly, in the *lec2* mutant the GUS activity was undetectable in the germinating seedlings (Figure 2G), as well as in 3-day-old plants (Figure 2H). This result demonstrates that the expression of the *ABI4* gene is fully dependent on the presence of the LEC2 regulator during germination and early seedling development.

**Figure 2.**
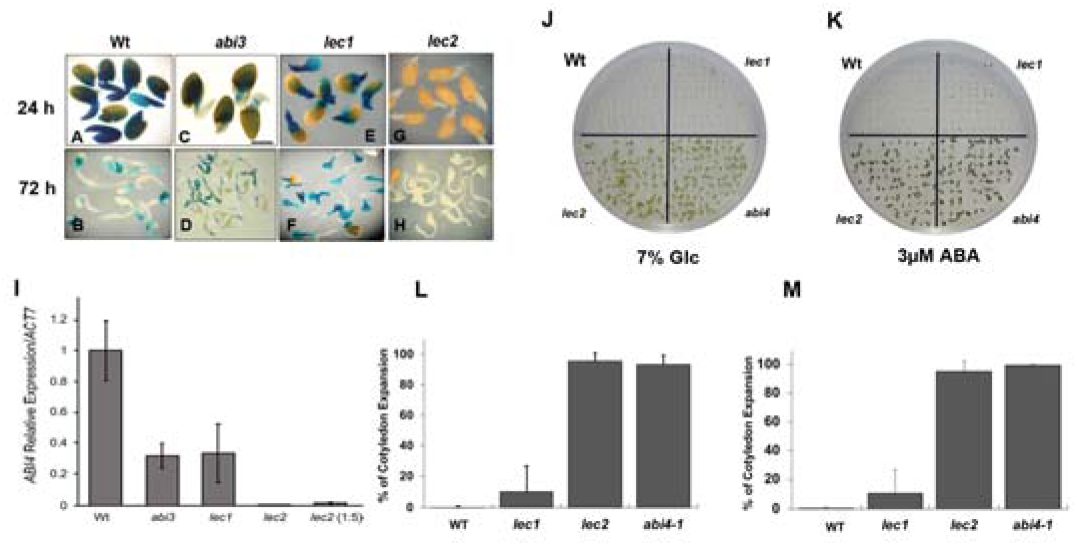
LEC2 is essential for the correct activation of ABI4 during early seedling development and for glucose and ABA signaling responses. Representative expression of *3KABI4:GUS* germinating seedlings at 24 h (A, C, E and G) and 72h (B, D, F and H) after transferred to germinating conditions for wild-type (Wt), *abi3*, *lec1* and *lec2* mutants. (I) Analysis by RT-qPCR of *ABI4* transcript levels from Wt, *abi3, lec1* and *lec2* mutant seedlings 24h after transference to germinating conditions. Transcript level using five times more *lec2* cDNA (1:5) is shown. Expression is reported relative to that of *Actin 7* (*ACT7*). Bars are means ±SE of triplicate biological experiments (each with n=2 technical replicates) and with P values p<0.05 between wild-type compared to the mutants (Student’s t test). Phenotypes of 14-day-old seedlings of Col-0 wild-type (Wt), *lec1*, *lec2* and *abi4* seedlings grown in the presence of media with 7% glucose (Glc) (J) or 3 μM ABA (L). (K) Percentage (%) of seedlings with expanded green cotyledons in the presence of 7% Glc (K) or 3 μM ABA (M). Error bars represent the SD of biological independent triplicate experiments.

To further explore the participation of LEC2 in the expression of *ABI4* during the early vegetative development, the transcript levels of *ABI4* were analyzed by quantitative real time PCR (RT-qPCR) in 24 h wild-type, *lec1 abi3* and *lec2* germinating seedlings, a time where the expression level of this gene is high in wild-type plants (Arroyo et al., 2003; Bossi et al., 2009). Our analysis showed a significant reduction in the accumulation of the endogenous *ABI4* transcript to approximately 60% in the *lec1* and *abi3* mutants, further supporting the role of these two transcription factors in the expression of this gene (Figure 2I). Moreover, similar to the GUS activity analysis, the *ABI4* endogenous transcript level in the *lec2* mutant was almost undetectable, showing at least 100-fold times lower expression than wild-type seedlings (Figure 2I), supporting the essential role of LEC2 for the expression of *ABI4*. Collectively our data demonstrate previously undescribed roles of the LEC1, ABI3 and, in particularly LEC2, in maintaining the expression levels of the *ABI4* gene during vegetative development.

### The *lec2* mutant is insensitive to ABA and sugar

It is known that ABI4 is required for proper ABA and sugar perception during early seedling development and its absence results in an ABA- (*abi*) and glucose- (Glc) (*gin*) insensitive phenotypes (Arenas-Huertero et al., 2000; Huijser et al., 2000; Finkelstein et al., 2011). To investigate whether the *ABI4* expression defects observed in the *lec1, abi3* and *lec2* seedlings affect the Glc and/or ABA sensitivity, we grew these mutants in the presence of 7% Glc or 3 μM ABA. These two conditions arrest greening and growth in wild-type seedlings, but not in *abi4* that behaves like the *abi* and *gin* mutants (Arenas-Huertero et al., 2000). We observed that in the presence of 7% Glc (Figure 2J and 2K) or 3 μM ABA (Figure 2L and 2M, more than 90% of the *lec1* seedlings became arrested similar to wild-type seedlings, demonstrating that the lower transcript levels of *ABI4* observed in this mutant do not result in *gin* or *abi* phenotypes, that is consistent with previous findings (Parcy et al., 1997). As previously reported (Dekkers et al., 2008), more than 90% of the *abi3* mutant seedlings displayed green cotyledons and continue growing in the presence of Glc or ABA (Figure S2), a phenotype similar to the *gin* and *abi* mutants. Also, this analysis showed that more than 90% of the *lec2* seedlings display clear *gin* and *abi* phenotypes, comparable to the *abi3* and the *abi4* seedlings, in the presence of 7% Glc (Figure 2J and 2K) or 3μM ABA (Figure 2L and 2M). These results confirm that the low levels of the *ABI4* transcript present in the *lec2* mutant seedlings results leads to alterations in the Glc and ABA sensitivity, further supporting a critical role of LEC2 in the regulation of *ABI4*.

### Proper *ABI4* expression depends on positive and negative *cis*-acting elements

The regulation of the *ABI4* expression by the LEC2 and ABI3 transcription factors could result from direct or indirect mechanisms. To further explore these possibilities, we analyzed the upstream regulatory region of the *ABI4* gene looking for putative ABI3 and LEC2 binding sites. These two transcription factors bind to RY DNA motifs or variants, containing the “CATG” core sequence (Braybrook et al., 2006; Swaminathan et al., 2008; Baud et al., 2016). Also, the presence of additional elements such as E- or G-boxes nearby can influence transcription factor binding (Abraham et al., 2016). The analysis of the 3 kb *ABI4* upstream sequence showed two sequences that fit the RY consensus elements. One of them, here referred to as RY1 (CATGCA), localizes −2467 bp upstream from the *ABI4* ATG and the other (RY2, GCATG) is at −2973 bp (Figure 3A). In addition, a canonical G-box (CACGTG) is present between the two RY motifs (Figure 3A). To analyze the possible participation of these RY elements in the transcriptional regulation of *ABI4*, we generated constructs containing consecutive deletions of the *ABI4* upstream sequence fused to the *GUS* reporter gene. The first deletion includes 2570 bp upstream from the *ABI4* ATG (2.5K*ABI4*) and lacks the RY2 and G-box elements (Figure 3A). The second deletion removed both RY elements as well as the G-box (2K*ABI4*) leaving 1990 bp of the upstream *ABI4* sequence (Figure 3A). Both deletions retained the ABI4 binding site (CE element), that localizes near the transcription initiation site. We generated transgenic plants carrying each deletion and the expression of GUS was analyzed in independent lines and compared to lines carrying a 3K fragment (3K*ABI4*:GUS) (Soderman et al., 2000; Bossi et al., 2009). In contrast to the 3K*ABI4*:GUS lines (Figure 3B–F), the GUS activity in the *2.5KABI4* and *2KABI4* deletion lines was undetectable in all stages of the developing seed (Figure 3G–P). These results are consistent with the RY and/or the G-box being essential *cis*-acting elements for the trans-activation of *ABI4* during embryo development. Intriguingly, this analysis also showed that the 2.5K*ABI4* deletion lines, ectopic GUS expression in the funicle and the valves of the siliques (Figure 3I and 3R) not present in the 3K*ABI4* (Figure 3B–F and 3Q) or the *2KABI4* (Figure 3L–P and 3S) lines. Furthermore, this ectopic expression extended to other vegetative tissues in the 2.5Kp*ABI4* lines (Figures 3 and S3) including the primary (Figure 3T) and rosette leaves (Figure 3U), the primary root (Figure 3T) and the flowers (Fig. 3V). All these are tissues where *ABI4* expression was never previously observed with the 3K*ABI4* transgene (Soderman et al., 2000; Bossi et al., 2009). This ectopic *ABI4* expression was exclusive of the 2.5K*ABI4* deletion and was not observed in the 2K*ABI4* transgenic lines (Figures 3S–W and S3). These results demonstrated that within the 500 bp between −3 kb and −2.5 kb of the *ABI4* upstream sequence there are essential *cis*-acting elements required to activate the expression of *ABI4* in the developing seeds and young seedlings and also for the repression of this gene in vegetative tissues.

**Figure 3.**
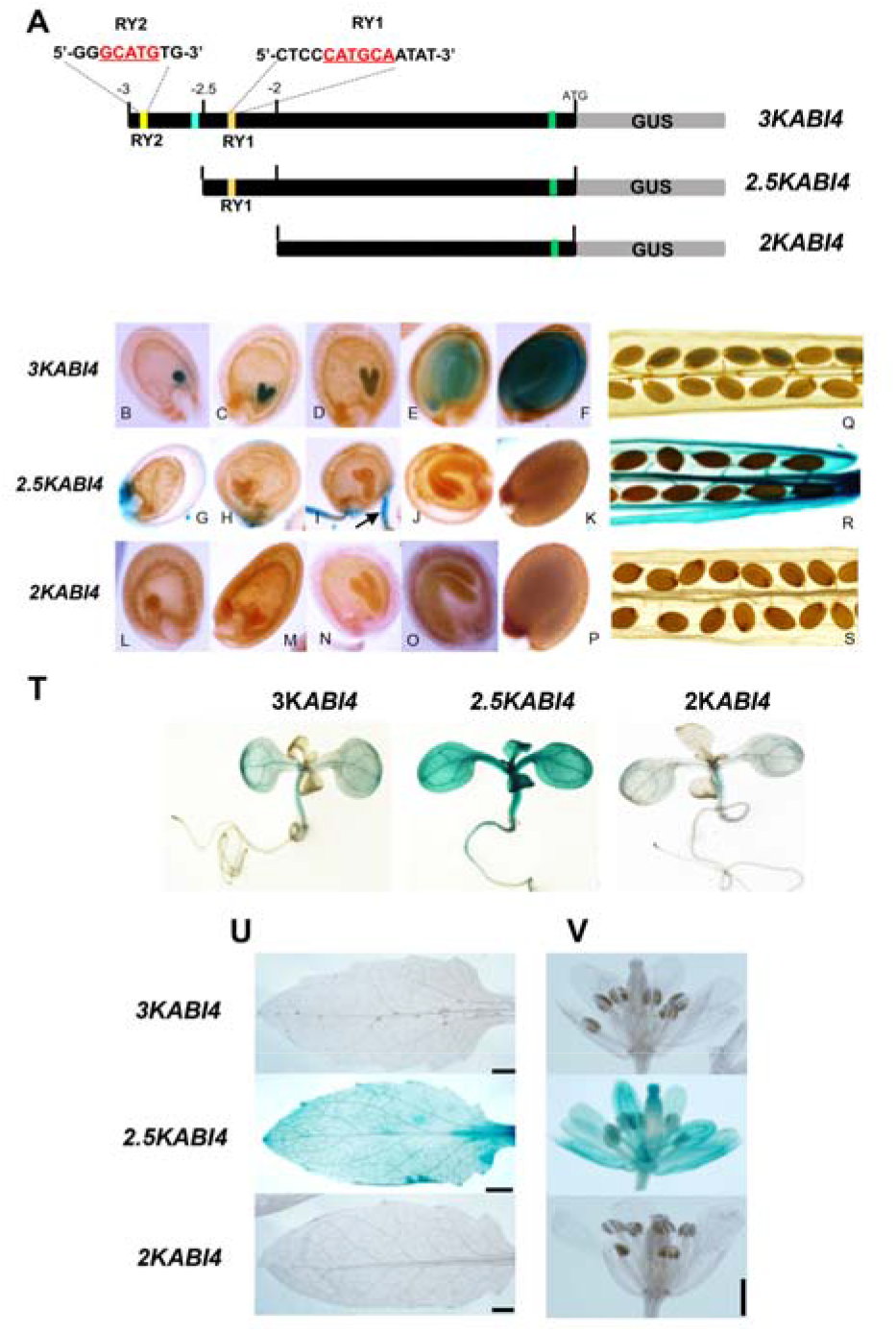
Analysis of the upstream regulatory region required for the *ABI4* gene expression. (A) Diagram of the upstream regulatory region of the *ABI4* gene showing the deletion fragments generated, marked in kb from the ATG. The location within the 3 kb upstream region of the two RY motifs and their corresponding sequences (yellow boxes), the putative G-box (blue box) and the CE element are indicated. Histochemical expression pattern of the GUS reporter in embryos at globular (B, G, L), heart (C, H, M), torpedo (D, I, N), bent cotyledon (E, J, O) and mature (F, K, P) stages and from siliques (Q, R, S), 14 day-old seedlings (T), rosette leaves (U) and flowers (V) tissues from representative transgenic lines expressing GUS from 3kb (*3KABI4*), 2.5 kb (*2.5KABI4*) and 2 kb (2*KABI4*) upstream sequences from the ATG of *ABI4*. The arrow points to maternal tissues. Scale bars: 500 μm.

### The RY motifs are essential for proper *ABI4* expression

To further dissect the function of the RY motifs in the activation and/or repression of the *ABI4* gene, we generated site-specific mutants in each element. We replaced six bases that included the core CATG sequence in the RY motifs with an AAATTT sequence using the 3K*ABI4*:GUS construct as template (Figure 4A) and generated transgenic lines for the single and double mutants. Interestingly, the lines containing mutations in the RY1 (mRY1) or in the RY2 (mRY2) resulted in undetectable GUS activity in the embryo seeds (Figure 4C and D and S4), compared to the 3K*ABI4* lines (Figure 4B). On the other hand, in these mutant lines we observed ectopic GUS expression in vegetative organs including leaves and flowers (Figure 4C and 4D). This GUS expression pattern correlates with the one observed in the 2.5K*ABI4* deletion lines (Figure 3). Finally, the GUS expression pattern of the lines containing both mutations (mRY1 RY2) was indistinguishable from the single mutants in the embryo and vegetative tissues (Figure 4E). Altogether, these results further demonstrate RY1 or RY2 motifs as essential *cis*-acting sites not only for the *ABI4* induction in the developing seed, but also for its repression in vegetative tissues and indicates that both RY elements are required for the correct expression of the *ABI4* gene.

**Figure 4.**
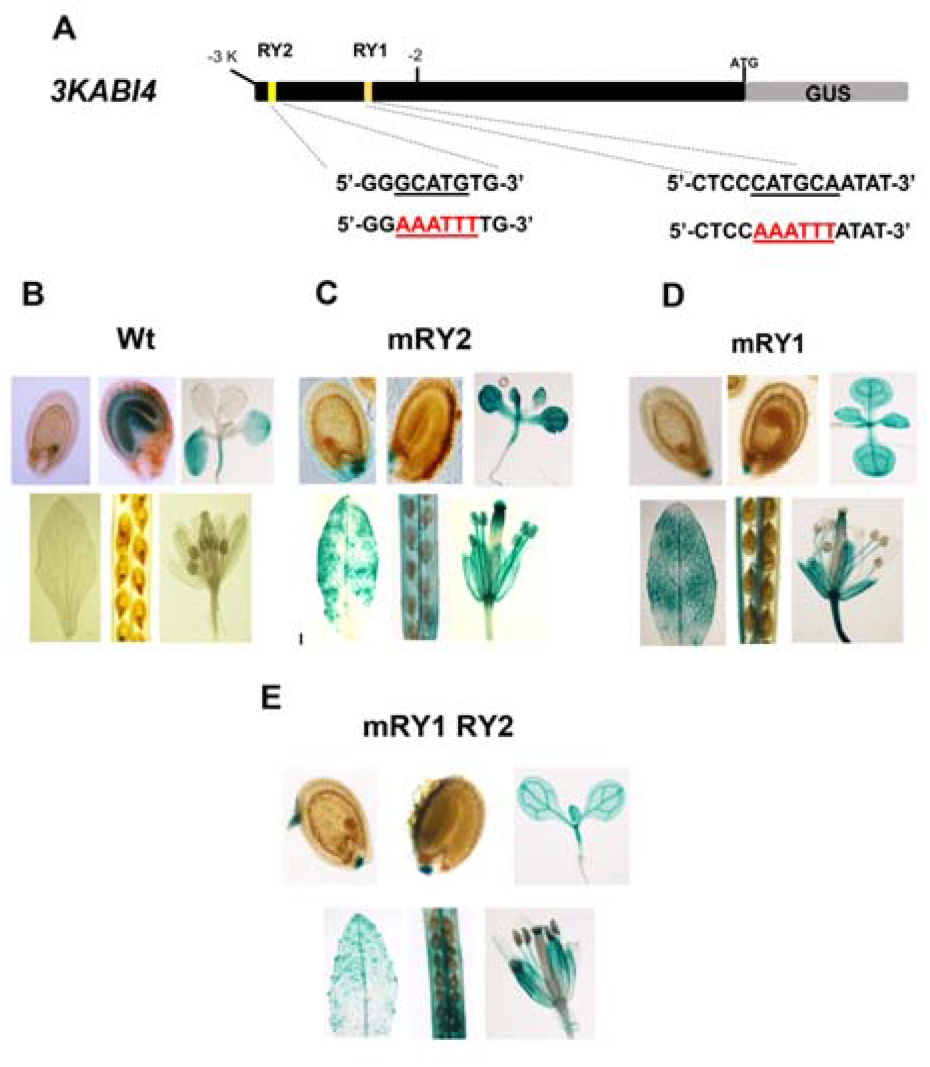
The RY motifs present in the regulatory region of *ABI4* are essential for the correct expression of the *ABI4* gene. (A) Diagram of the upstream region of *ABI4* showing the mutations generated in the RY elements. The changes introduced in each mutant construct are indicated in red compared to the original sequence. Histochemical expression of seeds at globular and bent cotyledon stages or in 14 day-old seedlings, rosette leaves, siliques and flowers from transgenic representative lines expressing GUS from the 3kb *ABI4* upstream sequence containing the (B) original RY motifs (Wt) or the site-specific RY mutations in the (C) RY2 (mRY2), (D) RY1 (mRY1) and (E) the double RY1 RY2 (mRY1 RY2) motifs.

### The expression of ABI4 is repressed in vegetative tissues by the HSI2/VAL1 and HSL1/VAL2 repressors

The RY motifs are known to be the binding site for the B3-domain regulators including HSI2/VAL1 and HSL1/VAL2 (HSI/VAL) factors, two proteins that mediate transcriptional repression of different genes through their interaction with chromatin-modifying proteins (Suzuki et al., 2007; Veerappan et al., 2014; Tao et al., 2019). Considering the ectopic expression observed for the *ABI4* gene in vegetative tissues when the RY elements were deleted or mutated, we reasoned that the HSI/VAL proteins were probably responsible for the repression of *ABI4* expression in vegetative tissues. To verify this hypothesis, we determined the expression level of *ABI4* by RT-qPCR in 10 day-old *hsi2 hsl1* loss-of-function mutant seedlings, because at this stage the *ABI4* transcript level is almost undetectable in wild-type plants (Tsukagoshi et al., 2007; Bossi et al., 2009). In agreement with our hypothesis, we observed a 100-fold accumulation of the *ABI4* transcript in the *hsi2 hsl1* mutant compared to wild-type seedling (Figure 5A). This result is consistent with the ability of HSI/VAL proteins to repress *ABI4* expression in vegetative tissues. This high *ABI4* transcript level in the *hsi2 hsl1* mutant also correlates with Glc- (Figure 5B) and ABA-hypersensitive (Figure 5D) phenotypes in these mutant seedlings, as previously reported for sucrose (Tsukagoshi et al., 2007). This result further supports the idea that high *ABI4* transcript level in the *hsi2 hsl1* mutant translates into higher ABI4 activity.

**Figure 5.**
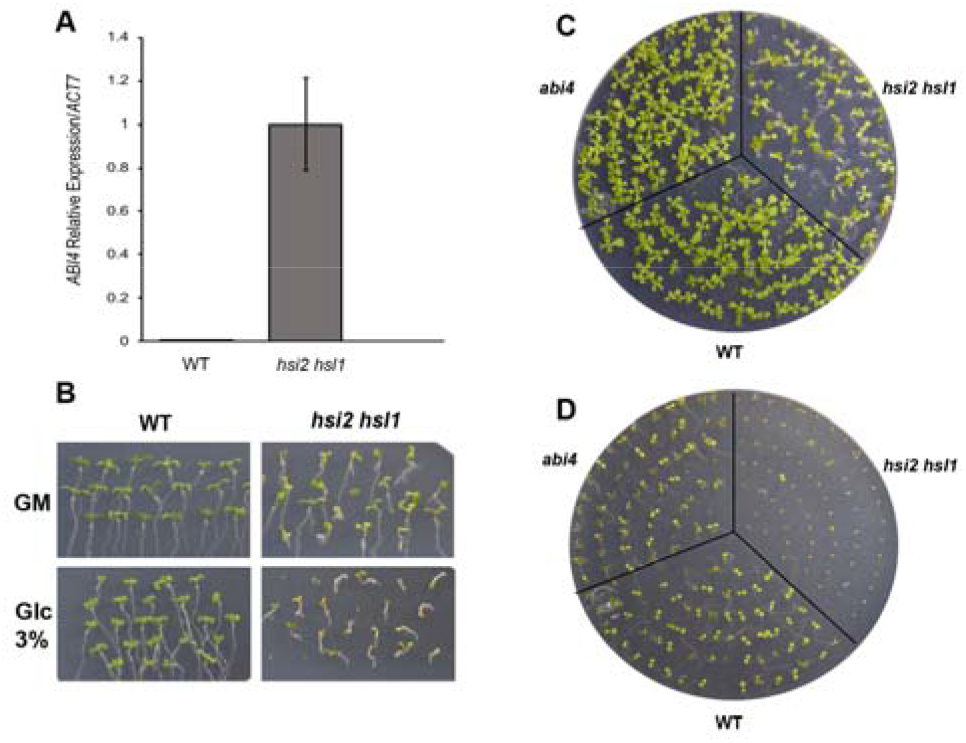
HSI2/VAL1 HSL1/VAL2 transcription factors are required for the correct repression of *ABI4*. (A) Analysis by RT-qPCR of the *ABI4* transcript accumulation in wild-type (Wt) or the *hsi2 hsl1* loss-of-function double mutant 10 day-old seedlings. *ABI4* expression is reported relative to that of *Actin7* (*ACT7*). Bars are means ±SE of triplicate biological experiments (each with n=2 technical replicates). (B) Phenotypes of 10 day-old Col 0 wild-type (Wt) and *hsi2 hsl1* mutant seedlings grown in the presence of 3% glucose (Glc). (C) Phenotypes from 15 day-old Col-0 wild-type (Wt), *hsi2 hsl1* and *abi4* mutants seedlings grown on GM media or (D) GM media in the presence of 0.5 μM ABA.

### LEC2 and HSI2/VAL1 bind to the same RY elements resulting in changes in chromatin accessibility

Altogether our new data demonstrates that LEC2 is an essential activator of *ABI4* expression, whereas ABI3 has an important, but not essential, contribution in its transcription. In addition, we provide unequivocal evidence that the HSI/VAL repressors are required for silencing *ABI4* gene expression in the vegetative tissues. Furthermore, our molecular analyses showed that the two RY motifs (RY1 and RY2) present in the *ABI4* upstream sequence are not only required for its activation by LEC2 and ABI3, but for HSI/VAL-dependent repression.

Since the LEC2, ABI3 and HSI/VAL proteins recognize the same *cis*-acting sequences, we reasoned that it was possible that these factors interact directly with these motifs. To investigate this possibility we mined available public genome wide chromatin immunoprecipitation coupled with high throughput sequencing (ChIPseq) data recently published for the LEC2 and HSI/VAL proteins (Wang et al., 2020; Yuan et al., 2021). The ChIPseq analysis for LEC2 was carried out in explants where LEC2 was induced by dexamethasone from the 35S::LEC2-GR-3X FLAG construct (Wang et al., 2020). Using this data, we corroborated a clear binding peak between −2K to −3K of the *ABI4* upstream sequence enriched in the LEC2-induced sample (DEX) compared to the control (Figure 6A). This region includes the two RY motifs that we showed are required for the *ABI4 trans*-activation in the developing seed (Figure 6). Thus, our data is fully consistent with the direct *trans*-activation of *ABI4* by LEC2. Moreover, a recent publication using ChIP-chip analysis of a p*ABI3*:ABI3-HA tagged transgene reported *ABI4* as an ABI3-associated gene, also supporting a direct interaction of ABI3 to the *ABI4* locus (Tian et al., 2020). Unfortunately, we were not able to identify the ABI3 binding region in the *ABI4* locus, but we hypothesize that the two RY elements identified here are most likely the binding sites also for ABI3 since no other sequences that fit the known binding element are found close to the *ABI4* gene.

**Figure 6.**
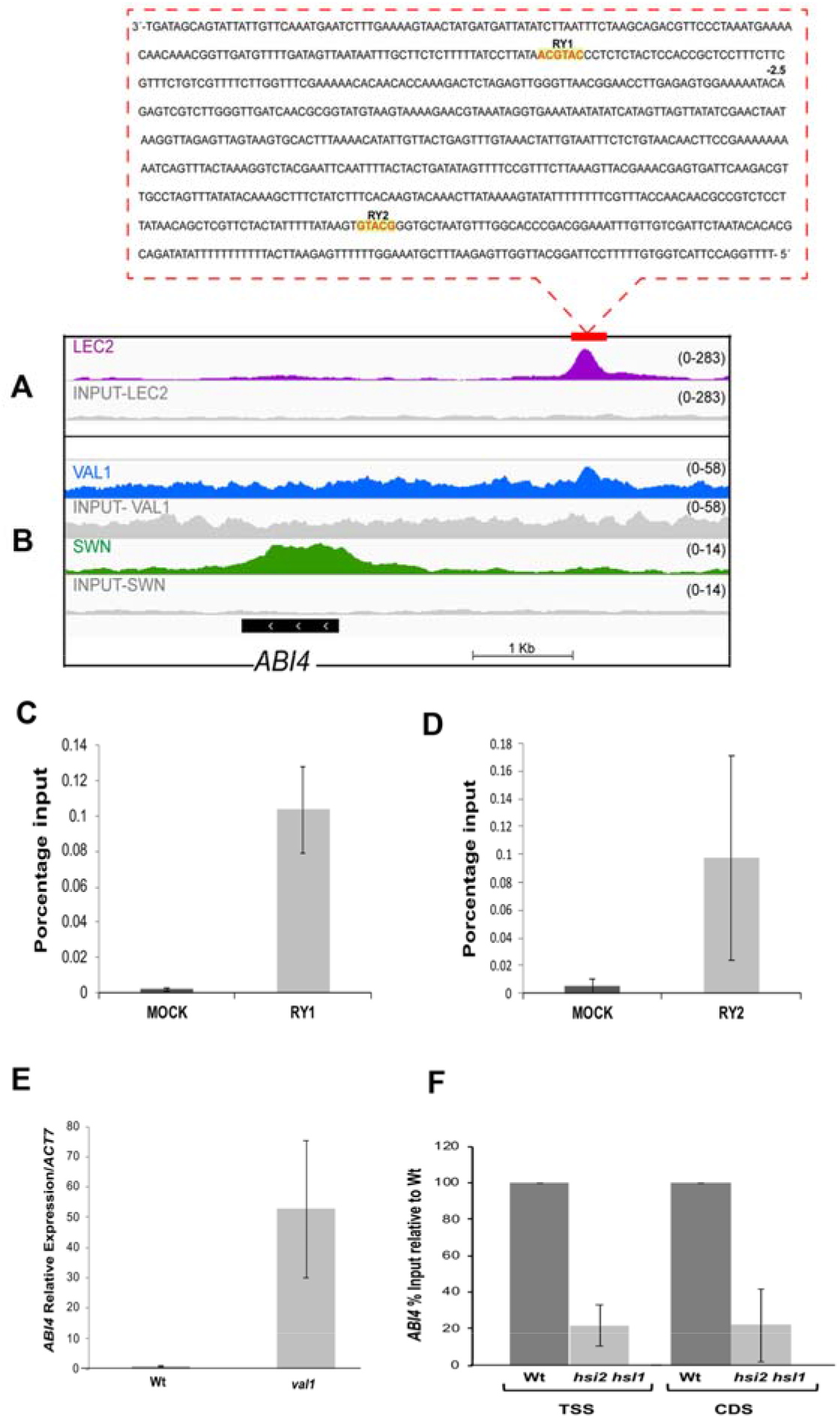
LEC2 and HIS/VAL factors bind to the *ABI4* upstream regulatory sequence. ChIP-seq signal for the *ABI4* loci for the (A) LEC2-GR-3xFLAG (Wang et al., 2020) and for the (B) VAL1-GFP and SWN-GFP (Yuan et al., 2021) DNA binding factors. The binding peaks in each case were detected and compared to the corresponding negative control. The location of the *ABI4* coding region is indicated by the black box. In the upper box the sequence included in the peaks that is common to the LEC2 and VAL1 regulators is shown and the location of the two RY elements is highlighted in yellow. Chromatin inmmunoprecipitation (ChIP) of the VAL1-HA protein binding in 10-day-old Col-0 (Wt) (dark grey) and VAL1-HA (light gray) seedlings along the *ABI4*. The qPCR from the immunoprecipitated sample was done using the pABI4-RY1-Fw/ and RY1 Chip qPCR Rv (C) or the pABI4-RY1RY2-Fw/ RY2 Chip qPCR Rv (D) primer pairs covering around the RY1 and RY2 elements. (E) Analysis by RT-qPCR of the *ABI4* transcript accumulation in wild-type (Wt) or the *val1* 10 day-old mutant seedlings. *ABI4* expression is reported relative to that of *Actin7* gene (*ACT7*). (F) ChIP analysis of the H3K27me3 in 10-day-old Col-0 (Wt) (dark grey) and *hsi2 hsl1* mutant (light gray) seedlings. The qPCR from the immunoprecipitated sample was done using the ABI4-ChIP-F and ABI4-ChIP-R primer pair covering the transcription initiation site of *ABI4* (TSS) and the primers ABI4-558Fw and ABI4-784Rv in the body of the ABI4 gene. The bars are the mean + SE of triplicate independent experiments (each with technical duplicates n = 2).

Finally, the potential interaction of the HSI2/VAL1 and/or HSL1/VAL2 proteins was analyzed using the ChIPseq data available from young seedlings expressing the VAL1-GFP or VAL2-GFP fusion proteins (Yuan et al., 2021). Using the published data, we observed an enrichment of a HSI2/VAL1 binding peak in the upstream region of *ABI4* (Figure 6B). However, due to a high basal background present in this study the interaction of the HSL1/VAL2 was not clear (data not shown). To further confirm this interaction chromatin immunoprecipitation (ChIP) analysis was performed using a Val1-HA transgenic line (Questa et al., 2016). From this analysis we confirmed an *in vivo* binding of VAL1 to the *ABI4* sequences containing the RY1 (Figure 6C) and RY2 (Figure 6D) motifs. Interestingly, the interacting peaks observed for the LEC2 and HSI2/VAL1 proteins in these independent studies overlap the region that includes the two RY *cis*-acting motifs, further supporting the role of these sequences as the binding site of these two proteins. To further address the role of the HSL1/VAL2 in the *ABI4* regulation we analyzed by qRT-PCR the expression of *ABI4* gene in the single *val1* mutant background. We observed that the absence of VAL1 alone resulted in a more than 50 times higher *ABI4* expression in comparison to the wild-type Col-0 background demonstrating that VAL2 cannot compensate the absence of VAL1. However, the upregulation observed in the *val1* mutant is lower than the one in *val1 val2* double mutant (more than 100X) in comparison to the wild type (Figure 5A). This result supports that VAL2 also participates in *ABI4* repression, at least in the absence of VAL1.

Histone modifications play an important function in the gene silencing mediated by HSI/VAL regulators as a result of the recruitment of the Polycomb-Repressive Complex 2 (PRC2) that catalyzes the tri-methylation of K27 for histone (3H3K27m3) deposition and transcription repression (Yuan et al., 2021). To determine if the chromatin status of the *ABI4* gene correlates with the presence or absence of the HSI/VAL regulators, we analyzed the levels of H3K27m3 mark by ChIP followed by qPCR around *ABI4* transcription initiation site in 14 day-old wild-type and *hsi2 hsl1* mutant seedlings. Our experimental data shows that in the *hsi2 hsl1* mutant the deposition of the H3K27me3 mark around the *ABI4* transcription initiation site and in the body of the gene was significantly reduced, with only 17% of the amount found in wild-type seedlings (Figure 6C). This result demonstrates that the absence of the HSI/VAL repressors significantly dilutes the H3K27m3 mark, and that is consistent with an active chromatin state and a high *ABI4* expression levels observed in this mutant (Figure 5A). Moreover, our data is also consistent with a high deposition of the SWN subunit, the Arabidopsis H3K27 methyl-transferases of PRC2 complex, that was in ChIP-seq studies (Yuan et al., 2021) across the entire of the *ABI4* gene in vegetative tissues (Figure 6B). Collectively these results support a direct regulation by LEC2, ABI3 and HSI2/VAL1 in the up-regulation of seed *ABI4* repression during vegetative development that is critical for the correct regulation of this transcription factor during plant development.

## DISCUSSION

Evidence has accumulated demonstrating that the transcription factor ABI4 is an integrator for diverse functions during plant development (Chandrasekaran et al., 2020). Consistent with the multifaceted role of ABI4, its accumulation and activity are strictly regulated at the transcriptional and post-translational levels (Chandrasekaran et al., 2020; Eisner et al., 2021; Zhou et al., 2021). *ABI4* displays a restricted spatio-temporal expression pattern during plant development that is also modulated by environmental stimuli, such as nutrients, hormones and abiotic stresses (Chandrasekaran et al., 2020). The correct transcriptional regulation of *ABI4* is critical for proper plant growth and stress responses, such as ABA and sugar perception (Finkelstein, 1994; Arenas-Huertero et al., 2000; Arroyo et al., 2003), and to avoid physiological harm due to its overexpression (Shkolnik-Inbar and Bar-Zvi, 2010; Shu et al., 2016b). Accordingly, several transcription factors act as negatively regulate *ABI4* gene expression, including various WRKYs, RAV1 and SCR proteins (Shang et al., 2010; Cui et al., 2012; Feng et al., 2014). In contrast, ABI4 itself stands as a central activator of its own expression during early seedling development (Bossi et al., 2009) and in this study we show that this auto-regulatory mechanism extends to all stages of the developing seed.

Consistent with previous reports, we confirm that *ABI4* is expressed since very early stages in embryogenesis (Soderman et al., 2000; Penfield et al., 2006) and its transcript is maintained at high levels during all the seed development, except for the dry seed. Some regulators of *ABI4* expression during the developing seed have been reported (Huang et al., 2016b), but the mechanisms involved in ensuring the correct spatio-temporal expression have not been identified. In this study, we demonstrate that the LAFLs regulators play distinctive roles in the spatial-temporal regulation of *ABI4* not only during embryogenesis but also during vegetative development, except for FUS3 which does not appear to have a major contribution. In contrast, our work provides clear evidence that the rest of the LAFL transcription factors have differential contributions on the *ABI4* gene expression.

An important contribution of this work is the demonstration that LEC2 and ABI3 are key regulators for the *ABI4* gene expression. Specifically, our evidence shows that LEC2 is an essential activator of *ABI4* transcription, not only in the developing seed but is also required during early seedling development (Figure 7). Interestingly, although LEC2 is expressed mainly during early seedling development (Santos-Mendoza et al., 2008; Baud et al., 2016; Boulard et al., 2017; Lepiniec et al., 2018) the absence of LEC2 results in undetectable *ABI4* expression in the germinating seedlings and in ABA- and Glc-insensitive phenotypes, demonstrating that LEC2 is one of the critical regulators of the *ABI4* expression. Likewise, we observed that ABI3 plays an important role regulating *ABI4* expression, but in contrast to LEC2, this factor is not absolutely required as a significant proportion of embryos accumulate normal levels of the *ABI4* transcript even in its absence. Moreover, in contrast to LEC2, ABI3 regulatory function is mostly restricted to developing seeds, further supporting the unique role of LEC2 in the regulation of ABI4 expression.

**Figure 7.**
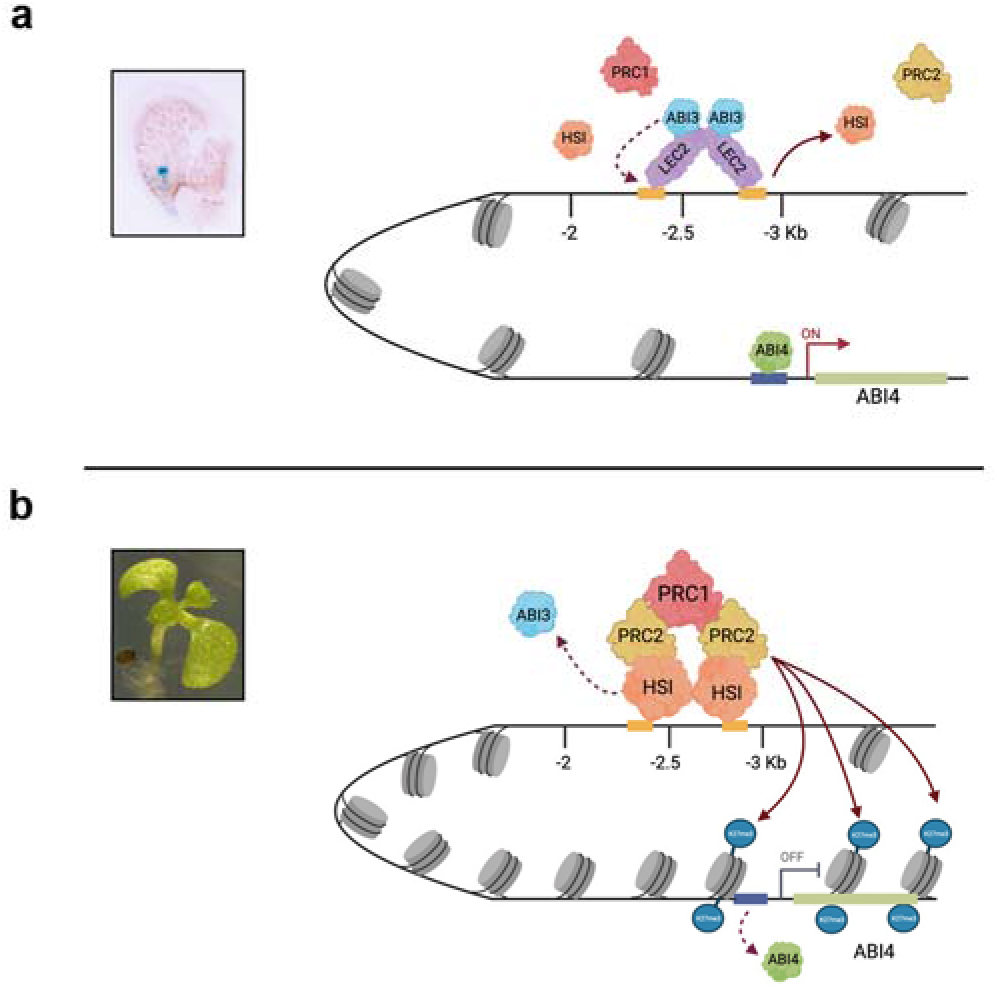
Model of the regulation of the *ABI4* gene expression by the LEC2, ABI3, HIS/VAL and ABI4 transcription factors. (A) The initial *ABI4* transcription activation occurs early in the of the developing seed (proembryo stage) by the initial activation by LEC2 dimer interacting with the two RY elements. This activation is critical for *ABI4* transcription initiation, probably establishing a permissive chromatin conformation schematized by a lower nucleosome number (grey cylinders) and no H3K27me3 mark. ABI3 binds to the same motifs, stabilizing LEC2 DNA as LEC2:ABI3 heterodimers or later by itself later once an open chromatin is established and the level of LEC2 decreases. The *de novo* translated ABI4 activates its expression during all developing seed stages. At these developmental stages the HSI/VAL repressors or the PRC2 complex do not interact with the *ABI4* locus due to the presence of LEC2 or ABI3. (B) During vegetative growth the absence of LEC2 and/or low ABI3 level together with the accumulation of HSI/VAL repressors allows these proteins to interact with the RY elements in the ABI4 locus. This interaction results in the accumulation of the PRC2 complex and probably the increase in the nucleosome number (grey cylinders) and the H3K27me3 mark; the transcriptional activity of the *ABI4* decreases and consequently the gene is silenced.

Additionally we observed a complex role for LEC1 in the regulation of *ABI4*. LEC1 acts as a negative regulator of *ABI4* during embryogenesis in the embryo suspensor tissue, but as an activator in young seedlings. Although the mechanism(s) involved in these antagonistic activities will require future analysis, we reasoned that they could result from indirect effects caused by the absence of LEC1. Because LEC1 is required for the suspensor specification (Lotan et al., 1998), and the ectopic *ABI4* expression observed in its absence might derive from the identity defects in this tissue. Also based on the previous observation that LEC1 activates the expression of *LEC2* and *ABI3* (To et al., 2006), the decrease of *ABI4* transcript levels observed during germination could be the consequence of a lower accumulation of these two transcription factors. Although has been shown that LEC1 and LEC2 or ABI3 can form complexes to regulate gene expression (Boulard et al., 2018), the regulation of *ABI4* is clearly distinct.

Importantly, we also provide new evidence that the correct modulation of *ABI4* expression relies not only on its activation in the embryo but also on its repression in vegetative tissues (Figure 7). We demonstrate that the repression of the *ABI4* in vegetative tissues is mediated by the HSI/VAL repressors; these repressors perform critical functions in the transition from embryo to seedling development (Suzuki et al., 2007; Veerappan et al., 2012). This negative regulation is critical in preventing the accumulation of *ABI4* in the growing plant, and causes ABA- and Glc-hypersensitivity as well as the downregulation of GA biosynthetic genes that are harmful for plant growth (Tsukagoshi et al., 2007; Shu et al., 2013).

The fact that the activation and the repression of the *ABI4* gene depends on the two RY elements located more than 2 kb upstream of this gene, supports that the observed regulation results from the binding of the LEC2, ABI3 and HSI2/VAL1transcription factors to the same two *cis*-acting motifs (Figure 7) and do not support that these regulators interact with other cis-acting elements, as was proposed for ABI3 (Tian et al., 2020). This is further corroborated by our ChIP analysis with VAL1, the published ChIP-seq data from the LEC2 and HSI2/VAL1 transcription factors (Tian et al., 2020; Wang et al., 2020; Yuan et al., 2021) and from the ChIP-Chip data of ABI3 (Tian et al., 2020). All of these studies identified *ABI4* as a direct target. This regulatory circuit could mediate an effective mechanism for fine-tuning of the correct spatial and temporal accumulation of ABI4 during plant development. Similar antagonism between the AFLs activation and HSI/VALs repression has been documented for other key genes involved in the ABA, GA and ethylene signaling (Braybrook et al., 2006; Stone et al., 2008), in the transition between embryogenesis and vegetative development, including the LAFLs (Wang and Perry, 2013; Jia et al., 2014) and in the regulation of flowering (*FLC*) (Tao et al., 2019).

Even though our study demonstrates that both LEC2 and ABI3 are important activators of *ABI4* gene expression, these two proteins clearly do not have the exact same function. As previously described, the expression of *ABI4* is fully dependent on LEC2, and this is not the case for ABI3. We hypothesize that the role of LEC2 resembles that of the pioneer activators (Zaret, 2018), that have the capacity to bind “*de novo”* to the *ABI4* locus early in embryogenesis promoting chromatin accessibility and activation of *ABI4* gene expression, and could potentially prevent the binding of HSI/VAL proteins (Figure 7). A similar pioneer transcription function has been documented for LEC1 (Tao et al., 2017).

Our results also suggest that the accessible chromatin status established by the initial activation by LEC2 can facilitate the recruitment of additional transcription regulators that may include ABI3 and the newly synthesized ABI4 (Figure 7). This recruitment could initiate the essential ABI4 feedback activation loop in the developing seeds and also during early seedling development (Bossi et al., 2009). Whether LEC2 alone or LEC2/ABI3 heterodimers could participate in this initial activation mechanism remains an open question for future analyses. The cooperation between LEC2 and ABI3 has been observed for the activation of the *OLE1* gene, where LEC2 and ABI3 bind to multiple RY motifs with a partial regulatory redundancy (Baud et al., 2016). This observation contrasts to our studies for *ABI4* where the two RY motifs are essential for positive and negative regulation. However, the fact that a proportion of the seeds have a normal *ABI4* expression pattern even in the absence of ABI3, supports the idea that LEC2 alone can fulfill the initial activation. It is possible that ABI3 participates in the *ABI4* transcriptional activation by either stabilizing the binding of LEC2 or by amplifying *ABI4* expression after its initial activation (Figure 7). This role might resemble that of the MYC factor that works as a general amplifier of transcription in human cells (Nie et al., 2020). The accessible chromatin status defined by LEC2 is likely maintained later in vegetative development and perhaps also in particular cell types by additional regulators in response to hormone and nutritional (Glc) levels (Tang et al., 2017), similar to what was reported for the *FLC* gene (Tao et al., 2019).

Later in development and probably as a result of the substantial decrease or total absence of LEC2 protein and the accumulation of the HSI/VAL repressors, chromatin remodeling of the *ABI4* locus will led to silence its expression (Figure 7); similarly to what has been reported for other seed maturation genes (Tsukagoshi et al., 2007; Shkolnik-Inbar and Bar-Zvi, 2011). In particularly, previous studies have demonstrated that HSI2/VAL1 and HSL1/VAL2 transcriptional repressors induce transcriptional silencing by promoting the trimethylation of the lysine 27 of histone 3 (H3K27m3) deposition as a result of their interaction with the Polycomb repressive complex 2 (PRC2) that is associated with gene silencing (Veerappan et al., 2014; Yuan et al., 2021). In support of this mechanism, we confirmed that *ABI4* repression correlates with high deposition levels of the H3K27m3 mark in the promotor region of *ABI4* and this depends on the presence of HSI2/VAL1 and HSL1/VAL2. This finding is similar to what has been described for other embryogenic genes including some of the LAFLs that promote the transition to vegetative development (Ogas et al., 1999). Based on the ChIP-seq data analyses for the HSI2/VAL1 and HSL1/VAL2 genes (Yuan et al., 2021), we observed that this repression correlates with an accumulation of the H3K27 methyl-transferase SWN of the PRC2 complex in the entire body of the *ABI4* gene (Figure 7).

In conclusion, this study describes a molecular mechanism that acts as a major spatio-temporal regulator of the *ABI4* transcription. This control mechanism results from the dynamic participation of multiple B3-type transcription factors that bind in the same *cis*-acting DNA elements to activate or repress *ABI4* gene expression. This precise control of the *ABI4* expression is mediated by changes in the chromatin state of the gene locus during specific moments of the plant life cycle.

## MATERIALS and METHODS

### Plant Material and Growth Conditions

Experiments were conducted in *Arabidopsis thaliana* L. Columbia-0 (Col-0) ecotype. Seedlings were grown on 1X GM media [Murashige and Skoog (MS) media with Gamborg vitamins (Phytotechology Laboratories, Shawnee Mission, KS), supplemented with 1% (w/v) sucrose, 0.5% MES and 0.8% (w/v) phytoagar] and stratified at 4° C for 4 days to break dormancy. Mature plants were grown in a 5:3:2 mixture of Peat moss 3 (Sunshine, Sun Gro Horticulture, Agawam, USA): vermiculite (Sun Gro Horticulture): perlite (Agrolita, Tlalnepantla, Mexico) containing 1.7 kg/m^3^ of Osmocote fertilizer (Everris, Geldermalsen, The Netherlands). Seedlings were grown in growth chambers (100 μmol m^−2^ s^−1^) and plants in walk-in chambers (80 μmol m^−2^ s^−1^) under 16:8h light:dark photoperiod at 22°C. To evaluate Glc or ABA sensitivity plants were grown on 1X MS medium containing 3% Glc or agar with 0.5 or 3 μM ABA and 0.5% MES.

The 3KABI4::GUS x *abi4* line was previously reported (Bossi et al., 2009). The 3KABI4::GUS transgene was introduced into the *lec1, lec2, fus3* and *abi3* mutant backgrounds by crossing the *pABI4*::GUS homozygous transgenic line (Soderman et al., 2000) with heterozygous plants of each mutant. The F2 progeny lines were selected for homozygocity of the transgene on 50 μg/mL kanamycin GM media and later the homozygous *lec1, lec2, fus3* and *abi3* mutant seeds were selected following the corresponding phenotypes. The homozygous *lec1, lec2, fus3* mutants lines carrying the pABI4::GUS transgene were maintained by germinating immature seeds on kanamycin media. The *val1* was obtained from the Arabidopsis Stock Center (SALK 088606C). The VAL1-HA transgenic line (Questa et al., 2016) and the *hsi2 hsl1* double mutant (Chen et al., 2020) were kindly provided by Drs. Caroline Dean (John Innes Center) and Allan Randy (Oklahoma State University).

### ABI4 promoter analysis

For the deletion analysis three constructs were generated, containing 3 Kb, 2.5 Kb and 2 Kb upstream of the start codon of *ABI4* using the pABI4 3Kb-FW, pABI4 2Kb-FW and pABI4 2.5Kb+attB1c FW 5’ oligonucleotides and the pABI4 −89 RV or RpABI4-89+B2c as 3’oligonucleotides (Supplemental Table 1). The fragments were introduced into the Gateway pMDC163 expression vector (Invitrogen, USA) to generate the corresponding transgenic lines (3K*ABI4,* 2.5K*ABI4* and 2K*ABI4*) through *Agrobacterium tumefaciens*-mediated transformation into the Col-0 ecotype (Clough and Bent, 1998). At least three independent homozygous lines were selected for each construct.

### Site directed mutagenesis of the RY elements

Mutants of the RY elements were generated by two step mutagenic PCR (Atanassov et al., 2009) using as template the 3 Kb fragment of the ABI4 upstream region and the oligonucleotides pABI4SphI-attB1 and ABI4-mRY1-Rv for the upstream region and the ABI4-mRY1-Fw and RpABI4-89+B2c for the downstream region (Table S1). For the RY2 mutation (mRY2) we used the oligonucleotides pABI4SphI-attB1 and ABI4-mRY2-Rv, and ABI4-mRY2-Fw and RpABI4-89+B2c (Table S1). The double RY1 and RY2 mutant was generated using the mRY2 fragment as templated and the mRY1 oligonucleotides. The reconstitution of the complete 3Kb *ABI4* fragments carrying the single or double RY mutations was obtained using the oligonucleotides pABI4SphI-attB1 and RpABI4-89+B2c (Table S1). The 3 Kb mutated fragments were introduced in the Gateway pMDC163 expression vector (Invitrogen, USA). mRY1, mRY2 or -mRY1RY2 transgenic plants were generated in the Col-0 ecotype (Clough and Bent, 1998). At least three independent homozygous transgenic lines were selected for each construct.

### Histochemical GUS Staining

Seedlings or plants were stained using the GUS histochemical assay (Jefferson, 1987). The tissues were vacuum infiltrated and incubated in GUS histochemical buffer (5mM of ferrous and ferricyanide) overnight. Plants were clarified according to a published protocol by (Malamy and Benfey, 1997). The tissues were semi-permanently mounted in a mix of 50 % glycerol and 2% DMSO and visualized using a stereoscopic (Nikon SMZ1500) or light (Nikon eclipse E600) microscopes.

### Expression Analysis

Total RNA was extracted from Col-0 seedlings using TRIzol (Thermo Fisher Scientific, Waltham, MA, USA) as recommended by the manufacturer. For RT-qPCR, RNA was treated with DNase (Promega, WI,USA) and cleaned following the instructions provided by the manufacturer (RNA clean & concentrator kit, Zymo Research). Complementary DNA (cDNA) was synthesized from 3μg of RNA using a M-MLV Reverse Transcriptase kit (Invitrogen, Carlsbad, CA, USA) and oligo dT. The RT-qPCR experiments were performed using Maxima SYBR Green/ROX qPCR Master Mix (Thermo Scientific, Baltics, UAB, Lithuania) on a Light Cycler 480 Roche. The oligonucleotides used in this analysis (ABI4-558Fw/ABI4-784Rv, for *ABI4* and ACT7-QPCR-F/ ACT7-QPCR-R, for *ACT7*) are listed in Table S1. Analyses were done with three independent experiments and technical duplicates were included in each case (n=2). The reference gene used in the qPCR analyses was *ACT7*.

### Chromatin Immunoprecipitation assays and *in silico* analyses

ChIP assays were conducted from 14 day old Col-0, *hsi2 hsl1* and VAL1-HA transgenic (Questa et al., 2016) seedlings grown on 1X GM media following the protocol previously reported (Johnson et al., 2002) with minor modifications. Tissue was cross-linked in fixation buffer (0.4M sucrose, 10 mM Tris-HCl [pH 8], 1 mM EDTA, 1mM PMSF and 1% formaldehyde) under vacuum. Samples were resuspended in lysis buffer (50mM HEPES [pH 7.5], 150mM NaCl, 1mM EDTA,1% Triton X-100, 0.1% deoxycholate, 0.1% SDS, 1 mM PMSF) and chromatin was sheared by sonication to approximately 500-100 bp fragments using the Bioruptor® sonicator (Diagenode, Belgium). Immunoprecipitation was done with the anti-H3K27me3 (Active-motif, Calsband, USA), anti-HA (Abcam, Cambridge, UK) or IgG antibodies (Invitrogen, USA). DNA-protein complexes were eluted from the Dynabeads (1% SDS and NaHCO_3_ 0.1M) and the crosslink was reverted with 5M NaCl. The RT-qPCR experiments were performed as previously described. The oligonucleotides used in these analyses were pABI4-RY1-Fw/ RY1 Chip qPCR Rv, pABI4-RY1RY2-Fw/ RY2 Chip qPCR Rv, ABI4-ChIP-F / ABI4-ChIP-R (Table S1).

For the analyses of the ChIP-seq, the raw data were downloaded from Gene Expression Omnibusunder accession numbers GSE119715 and GSE159499 (Yuan et al., 2021) and from Beijing Institute of Genomics Data Center, BioProject PRJCA002620 (Wang et al., 2020). Reads were aligned using Bowtie2 v2.3.4.3 (Langmead and Salzberg, 2012) to the *Arabidopsis* genome (TAIR10). The resulting SAM file containing mapped reads were converted to BAM format, sorted, and indexed using Samtools v1.9 (Li et al., 2009). Duplicated reads were removed using Picard tools (Picard Toolkit, 2019). Only perfectly and uniquely mapped reads were retained for further analysis. To normalize and visualize the datasets, the BAM files were converted to bigwig using bamCoverage provided by deepTools v3.1.2 (Ramirez et al., 2014). Finally, the bigwig files were visualized in the Integrated Genome Browser (IGV).

## AUTHOR’S CONTRIBUTIONS

PL, EC, MSM and AH designed the experiments; AH, EC, MSM, KAA, ADR and MU conducted the experiments; MU-A performed the bioinformatics analyses; MZ and PL analyzed the data; MZ and PL wrote/edited the paper. PL prepared figures.

## ACKNOWLEDGEMENTS

We are very grateful to Drs. Caroline Dean (John Innes Center) for kindly sharing the VAL1-HA transgenic line and Allan Randy (Oklahoma State University) for sharing the *hsi2 hsl1* double mutant. We thank Dr. Jen Sheen, Dr. Gabriela Toledo-Ortíz and Dr. Josefat Gregorio for their comments and suggestions and Dr. Kenneth Luehrsen for editorial suggestions. The IBT computer facility for access to the computer cluster. The authors declare no competing interests.

## Figure legends

**Supplemental Figure 1. Expression of *pABI4*:GUS in germinating seedlings.** GUS activity of *pABI4*:GUS transgene in the segregating heterozygous *lec2* (+/-) germinating seedlings. GUS activity was detected in three out of four heterozygous segregating plants.

**Supplemental Figure 2. *abi3* mutant displays a glucose- and ABA-insensitive seedling phenotype.** Phenotype of 10 day-old seedlings of Col-0 wild-type (Wt), *abi3* and *abi4* mutants in the presence of media containing 7% glucose (A) or 3μM ABA (B).

**Supplemental Figure 3. Expression of *pABI4*:GUS in germinating seedlings.** GUS histochemical activity in 14-day-old seedlings from independent transgenic lines carrying 3 kb, 2.5 kb and 2 kb of the *ABI4* regulatory region fused to the GUS reporter gene.

**Supplemental Figure 4. Expression of *pABI4*:GUS carrying mutation in the RY elements.** GUS histochemical activity in 14-day-old seedlings from independent transgenic lines carrying mutations in the RY1 (mRY1), RY2 (mRY2) and the double mRY1 and RY2 (mRY1 RY2) elements in the *ABI4* regulatory region fused to the GUS reporter.

